# *Cabbage1* and *Cabbage2* encode CYP90A orthologs that function redundantly to regulate plant development in maize

**DOI:** 10.64898/2026.11.12.766257

**Authors:** Brian Zebosi, John Ssengo, Erica Unger-Wallace, Norman B. Best, Erik Vollbrecht

## Abstract

Ideal plant architecture improves grain yield per unit area in maize fields by promoting upright leaves which support high planting densities, improving light interception and enhancing photosynthetic efficiency. Shoot architecture is defined by plant height, leaf morphology, and branching pattern, which are regulated by a complex network of plant hormones and environmental factors. Brassinosteroids (BRs) are among the classic plant hormones that regulate plant architecture. Here, we investigated the functions of *Cabbage* (*Cbg*) genes in maize which are orthologs of Arabidopsis *CYP90A1/CPD* and encode CYP90A, a key BR biosynthetic enzyme. Using reverse genetics, we identified and characterized *cbg1* and *cbg2* mutant alleles recovered from UniformMu transposon stocks. Our genetic mutant analysis indicated that *Cabbage* genes function collectively; each single mutant (*cbg1* and *cbg2*) was indistinguishable from its wild-type siblings. In contrast, *cbg1*;*cbg2* double mutants exhibited severe developmental defects including dwarf stature, fewer tassel branches, and smaller, erect leaves. These defects lead to shoots with tightly wrapped leaves forming a cabbage-like head that restricts inflorescence (tassel and ear) emergence and results in loss of fertility. Consistent with impaired BR biosynthesis, *cbg1;cbg2* double mutants had significantly reduced levels of downstream BR metabolites. Treatment of wild-type and single mutant germinating seedlings with the BR biosynthetic inhibitor, Propiconazole (PCZ), phenocopied untreated *cbg1*;*cbg2* double mutants, particularly in mesocotyl elongation. Supplementation with exogenous brassinolide (BL) alone was not adequate to restore mesocotyl elongation in the double mutants. These mutant phenotypes are consistent with impaired BR biosynthesis and demonstrate that *Cabbage* genes (*Cbg1* and *Cbg2*) function redundantly to regulate plant development. Overall, these findings on *Cabbage* genes highlight the potential of BRs for crop improvement.

## Introduction

Brassinosteroids (BR) are a major class of plant steroid hormones that regulate plant growth and development, as well as responses to biotic and abiotic stresses (Nolan et al. 2020; Zebosi et al. 2024). Genetic studies primarily in Arabidopsis (*Arabidopsis thaliana*) have shown that mutations in BR biosynthetic and signaling pathways result in dwarf to semi-dwarf plants with reduced fertility (Clouse 2002; Vriet et al. 2013). In monocots including maize (*Zea mays*) and rice (*Oryza sativa*), BR-related mutants similarly exhibit semi-dwarfism with erect leaves and altered reproductive organ development, highlighting conserved roles of BRs across species (Hong et al. 2005; Hartwig et al. 2011; Best et al. 2016; Zhan et al. 2022). Notably, mutant alleles with reduced BR activity and less severe pleiotropic effects produce semi-dwarf plants with erect leaves, agronomic traits that can enhance grain yield by improving light penetration under high planting densities (Sakamoto et al. 2006). For example, the maize quantitative trait loci *Upright Plant Architecture* (*UPA1* and *UPA2*), which map to the *brassinosteroid C-6 oxidase* (*brd1*) and 3-domain transcription factor gene, *ZmRAVL1*, respectively, regulate leaf angle through the BR pathway (Tian et al. 2019). Introgression of these alleles into maize hybrids produced plants with more erect leaves and increased grain yield under dense planting conditions, demonstrating the plausibility of manipulating the BR pathway for crop improvement (Tian et al. 2019).

In dicot plants, brassinolide (BL) is the most bio-active BR synthesized directly from Castasterone (CS), while CS is interpreted to be the bio-active BR in some monocots (Kim et al. 2008). These end products of the BR biosynthetic pathway are produced downstream of the phytosterol pathway from campesterol (CR), through parallel series of reduction, oxidation, and hydroxylation reactions catalyzed by a steroid 5α-reductase and several cytochrome P450 (CYP450) monooxygenases of the CYP90A, CYP90B, CYP90C, CYP90D, CYP85A, and CYP724B families (Fujioka and Yokota 2003; Ohnishi et al. 2006; Ohnishi 2018; Zebosi et al. 2024). Functional characterization of these enzymes through mutant analysis in Arabidopsis and rice has established their roles in plant development and stress responses (Clouse 2002). In particular, CYP90A, also dubbed CONSTITUTIVE PHOTOMORPHOGENIC DWARF (CPD), functions very early in the BR pathway as a C-3 oxidase that catalyzes the conversion of CR and its hydroxylated derivatives into corresponding 4-en-3-one intermediates (Ohnishi et al. 2012; Zebosi 2024).

While its enzymatic activity appears conserved, gene copy number of CYP90A/CPD varies across plant species. Arabidopsis contains a single gene (*AtCPD*). By contrast rice contains two paralogs (*OsCPD1* and *OsCPD2*) due to genome duplication, and the double mutant (*oscpd1*;*oscpd2*) exhibits severe growth defects, including dwarfism and upright leaves (Zhan et al. 2022). Similarly, the maize genome has two *CYP90A* homologs, which we have named *Cabbage1* (*Cbg1*) and *Cabbage2* (*Cbg2*) (Zebosi et al. 2024). To investigate the functions of CYP90As in maize, we characterized recessive mutants of *cbg1* and *cbg2* identified from the *UniformMu* transposon population (McCarty et al. 2005). Single mutants were phenotypically indistinguishable from their wild-type siblings. On the other hand, *cbg1*;*cbg2* double mutants exhibited severe developmental defects, including dwarf plant stature, fewer tassel branches, and smaller and erect leaves. BR metabolite profiling revealed reduced BR downstream intermediates in *cbg1;cbg2* double mutants and agonist/antagonist treatments were further consistent with disrupted BR biosynthesis. Our results suggest that *Cbg1* and *Cbg2* function reductantly in BR biosynthesis and regulate plant growth and development.

## Results

### Identification of *cbg1* and *cbg2* single mutants and generation of *cbg1;cbg2* double mutants

Cytochrome P450 monooxygenases of the CYP90A/CPD family encode brassinosteroid biosynthetic enzymes that play vital roles in plant growth and development (Ohnishi et al. 2012; Zhan et al. 2022). To determine the evolutionary relationships of CYP90A/CPD related genes across land plants, we used the Arabidopsis CYP90A (*AtCPD*) protein sequence to perform BLAST searches (Altschul et al. 1990; Camacho et al. 2009) against multiple plant genomes. We retrieved amino acid sequences with high similarity to CYP90A from Phytozome (Goodstein et al. 2012) and NCBI (NCBI Resource Coordinators 2014). The phylogenetic analysis included Arabidopsis, maize, rice, Brachypodium (*Brachypodium distachyon*), Medicago (*Medicago truncatula*), Sorghum (*Sorghum bicolor*), Setaria (*Setaria viridis*), soybean (*Glycine max*), tomato (*Solanum lycopersicum*), and Physcomitrella (*Physcomitrella patens*) as an out-group. Arabidopsis, tomato, Sorghum, and Brachypodium contain a single *CYP90A/CPD* gene, whereas the other surveyed species contain multiple *CYP90A/CPD* homologs consistent with several lineage-specific genome duplication events (**Figure 1A**, (Zebosi et al. 2024)).

**Figure 1:**
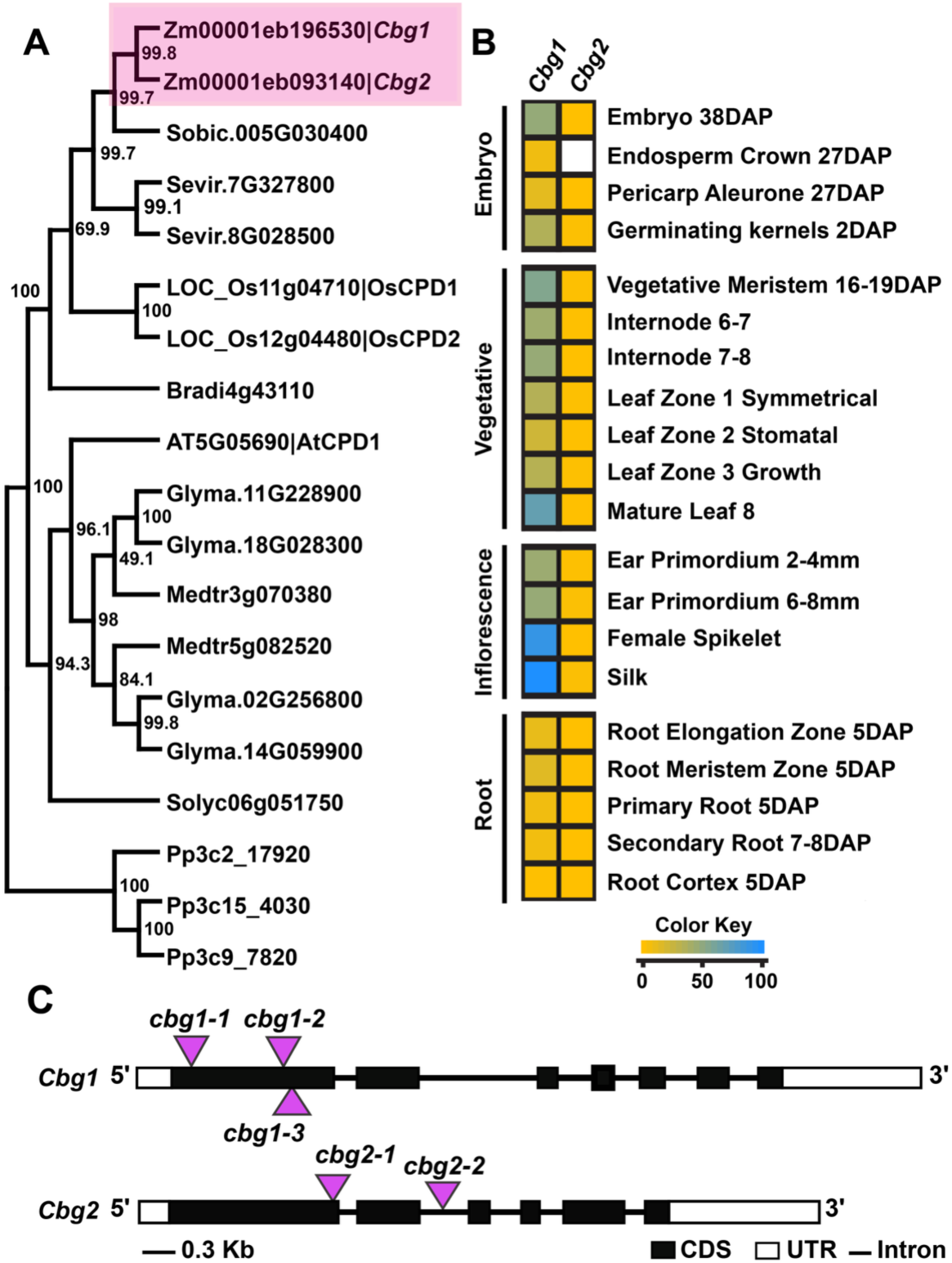
Phylogenetic analysis and identification of *CBG1* and *CBG2 mutator* insertional alleles. **A.)** Phylogenetic tree of CYP90A amino acid sequences from *Arabidopsis*, Brachypodium, Maize, Setaria, Sorghum bicolor, Medicago, soybean, rice, tomato and Physcomitrella, generated using the maximum likelihood method. Bootstrap values from 1,000 replications are displayed at the branch points. **B).** mRNA expression levels of *Cbg1* and *Cbg2* across several tissues based on Walley et al. (2016). **C).** Gene structure with Uniform *Mutator* transposon insertions (denoted by triangle) and corresponding allele designations.

The maize genome contains two duplicate CYP90A homologs, *Cabbage1* (*Cbg1*) and *Cabbage2* (*Cbg2*), which share approximately 60% and 58% amino acid identity with *AtCPD1*, respectively. *Cbg1* and *Cbg2* reside on syntenic regions of chromosomes 2 and 4, respectively, and likely arose from a recent whole-genome duplication (Schnable et al. 2011; Hughes et al. 2014). Based on subgenome assignments, *Cbg1* and *Cbg2* belong to the maize-1 and maize-2 subgenomes, respectively (Schnable et al. 2011). The two genes share 86% amino acid identity and 86.4% nucleotide identity, suggesting potential functional redundancy (Danisman et al. 2013; Panchy et al. 2016). Reanalysis of publicly available RNA-seq data (Walley et al. 2016), showed that *Cbg1 is* more highly expressed than *Cbg2* across most tissues (**Figure 1B**), suggesting unequal contribution to plant development. This expression difference is consistent with biased fractionation following whole-genome duplication, in which genes retained in the maize-1 subgenome are generally more highly expressed than their maize-2 homeologs (Schnable et al. 2011).

To test the functional relationship between *Cbg1* and *Cbg2*, we analyzed putative *Mutator* (*Mu*) transposon-insertion alleles from the UniformMu transposon population, which is in a uniform, inbred W22 genetic background (McCarty et al. 2005; Settles et al. 2007; Woodhouse et al. 2025). We recovered three independent *cbg1* alleles *(cbg1-1*, *cbg1-2*, and *cbg1-3*) and two *cbg2* alleles (*cbg2-1* and *cbg2-2*) for functional characterization (**Figure 1C**). PCR genotyping confirmed that all *cbg1* insertions reside in coding regions of exon 1, while *cbg2-1* and *cbg2-2* are located in the coding region of exon 1 and in intron 2, respectively. Based on their insertion sites, these alleles are predicted to disrupt gene function through premature stop codons or aberrant splicing (Simon and Starlinger 1987; Feiz et al. 2024).

When the insertion alleles were made homozygous in segregating populations, none of the single mutants (*cbg1-1*, *cbg1-2*, *cbg1-3*, *cbg2-1*, or *cbg2-2*) exhibited apparent developmental defects. Single mutants were indistinguishable from their wild-type siblings under both field and greenhouse conditions (**Supplemental Figures 1, 2, 3 and 4**). The absence of obvious developmental defects across multiple independent alleles of both genes suggests that *Cbg1* and *Cbg2* function redundantly during plant growth and development.

### *CBG1* and *CBG2* function redundantly to regulate plant growth and development

To test the hypothesized genetic redundancy, we generated four independent double mutant populations (*cbg1-1*;*cbg2-1*, *cbg1-2*;*cbg2-1*, *cbg1-3*;*cbg2-1* and *cbg1-1*;*cbg2-2*) by intercrossing pairs of single mutants and selfing the resulting F1 plants to produce four F2 populations. We recovered 11/130, 9/95, 14/135, and 18/185 double mutants from the respective populations, consistent with the expected 1:16 Mendelian segregation ratios (χ2, P > 0.05 in all populations). To address the possibility that altered phenotypes (described below) resulted from genetically linked background mutation, we also generated allelic compound heterozygous double mutants at both loci (*cbg1-1/cbg1-2;cbg2-1, cbg1-1/cbg1-3;cbg2-1, cbg1-2/cbg1-3;cbg2-1*, and *cbg1-1;cbg2-2/cbg2-1*). All compound heterozygous combinations failed to complement the double mutant phenotype, confirming that the double mutant phenotype is the result of independent mutations in *Cbg1* and *Cbg2* and indicating they have redundant function.

Unlike single mutants, double mutant plants from all four populations (*cbg1-1;cbg2-1*, *cbg1-2;cbg2-1*, *cbg1-3;cbg2-1*, and *cbg1-1;cbg2-2*) displayed developmental defects as early as 7-10 days after planting, with dwarfism as the earliest and most obvious phenotype. At maturity, the double mutants ranged from dwarf to semi-dwarf, reaching 12-55% of the wild-type height. The reduced plant stature was accompanied by compressed internodes, and the severity of the dwarf phenotype varied among allelic combinations (**Figure 2; Supplemental Figures 1 and 2)**. Double mutants with the *cbg2-1* allele were consistently more severely dwarfed than those carrying the *cbg2-2* allele (**Figure 2A; Supplemental Figure 3**), such that *cbg2-2* behaved as a weaker allele than *cbg2-1*.

**Figure 2:**
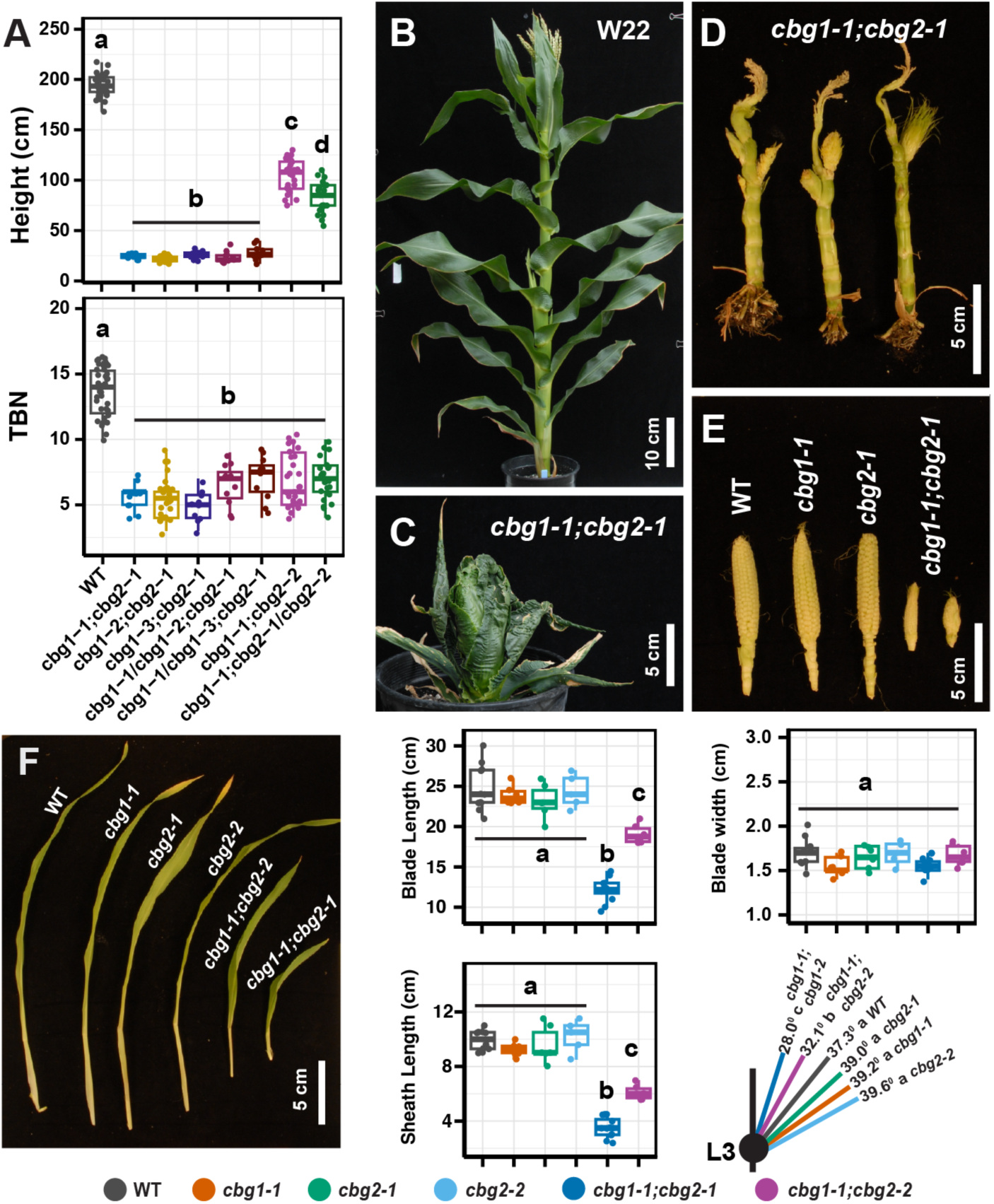
**Effect of *cbg1;cbg2* on plant height, tassel branching and leaf morphology**. **A).** Mature wild-type plant (W22). **B-C).** Mature *cbg1-1;cbg2-1* double mutant plants showing reduced plant stature and compressed internodes. Plants are shown with leaves intact in panel **B** and with leaves removed to expose the internodes in panel **C**. **D**). Ear size across genotypes at 3 months of age. **E).** Plant height and tassel branch number (TBN) at maturity across different *cbg1;cbg2* mutant allele combinations. Sample sizes were n=80, 20, 48, 22, 20, 20, 56 and 50, respectively. **F).** Blade length, blade width, sheath length (cm), and leaf angle of leaf 3 across genotypes; WT (n=11), *cbg1-1* (n=6), *cbg2-1* (n=6), *cbg2-2* (n=5), *cbg1-1;cbg2-1* (n=12), and *cbg1-1;cbg2-2* (n=6). Colored circles indicate the genotype represented in the graphs. Different lowercase letters indicate significant differences based on one-way ANOVA followed by Sidak multiple comparison tests (P < 0.05)

In addition to reduced plant height, *cbg1;cbg2* double mutants exhibited reproductive defects, producing smaller ears (**Figure 2E**) and 42–50% fewer tassel branches than their wild-type siblings and single mutants (**P < 0.05**; **Figure 2E; Supplemental Figure 1)**. Double mutants with the *cbg2-1* allele formed extremely twisted leaf whorls that developed into a compact, cabbage-like head (**Figure 2C**) that physically hindered emergence of reproductive organs (tassel and ear) (**Figure 2C**). These leaf twisting defects became evident around leaf 8 and persisted onward. In the context of these morphological defects, tassels were ultimately rendered infertile but produced aberrant anthers that failed to shed pollen, and ears were extremely reduced in size (**Figure 2E**) but produced silks, suggesting that female fertility may be retained if manually pollinated. These pleiotropic defects resemble those reported for the maize *brd1* mutant (Makarevitch et al. 2012), except that no tassel feminization was observed in *cbg* mutants. Brassinosteroid deficient mutants have not been previously described in the W22 genetic background (Hartwig et al. 2011; Makarevitch et al. 2012; Best et al. 2016), so the absence of the tassel feminization phenotype may be due to genetic background effects.

Double mutants also exhibited several defects in leaf development. In addition to pronounced leaf waviness and twisting, they produced shorter leaf blades and sheaths than their wild-type and single mutant siblings as quantified in young, unencumbered leaves (P < 0.05) (**Figure 2F; Supplemental Figure 4A**). On the contrary, blade width was unchanged in young leaves (Leaf 3) but significantly increased in older leaves (Leaf 6) (**Figure 2F; Supplemental Figure 4A**). Reduced blade length, rather than blade width, is a hallmark phenotype of BR-deficient and BR-insensitive mutants, consistent with the established role of BR in promoting longitudinal cell expansion (Hartwig et al. 2011; Li et al. 2013). Double mutants also displayed aberrant auricle/ligule boundaries, characterized by consistently enlarged auricles and a poorly defined auricle/blade boundary (data not shown). Consistent with these alterations, leaves of double mutants were more erect, with leaf angles about 10-13 degrees more upright than those of wild-type sibling and single mutants (**Figure 2F; Supplemental Figure 4B**). Leaf angle measurements were taken before leaf 8 emerged and prior to visible evidence of the cabbage head-like structure (**Supplemental Figure 3**).

### CYP90A1 mutants show reduced accumulation of BR biosynthesis intermediates

To test the hypothesis that *Cbg1* and *Cbg2* function in BR biosynthesis, we measured levels of select phytosterols and BR intermediates in vegetative shoot tissue of wild-type and sibling single and double mutant seedlings. Across all genotypes, levels of bulk sterols and stigmasterol were not significantly different between the genetic variants (**Supplemental Figure 5**). Similarly, levels of the high-abundance BR precursor Campesterol (CR), the first compound in the BR pathway, and of its immediate precursor the relatively low abundance phytosterol 24-methylene cholesterol were both not significantly between the mutants and wild type (**Figure 3A and Supplemental Figure 5**). In contrast, two BR metabolites downstream of CYP90A1, campestanol and its downstream intermediate 6-oxocampestanol, each accumulated to significantly reduced levels in the *cbg1-1* and *cbg2-1* single mutants. Each single mutant showed a similar reduction for each metabolite. In the *cbg1-1*;*cbg2-1* double mutant a further reduction was significant for campestanol levels. 6-oxocampestanol levels were reduced in the double mutant, as compared to the level in either single mutant (**Figure 3A**), but the reduction was not significant by a multiple test correcting *p*-value. Interestingly, campestanol and 6-oxocampestanol levels were equally affected in each (*cbg1-1* and *cbg2-1*) single mutant, even though in wild-type shoots *Cbg1* transcripts are expressed more highly than *Cbg2* transcripts (**Figure 1B**). These metabolite measurement results confirm that *Cbg1* and *Cbg2* act in BR biosynthesis in maize and that reduced BR levels are likely the cause of altered plant architecture phenotypes in *cbg1-1*;*cbg2-1* double mutants.

**Figure 3:**
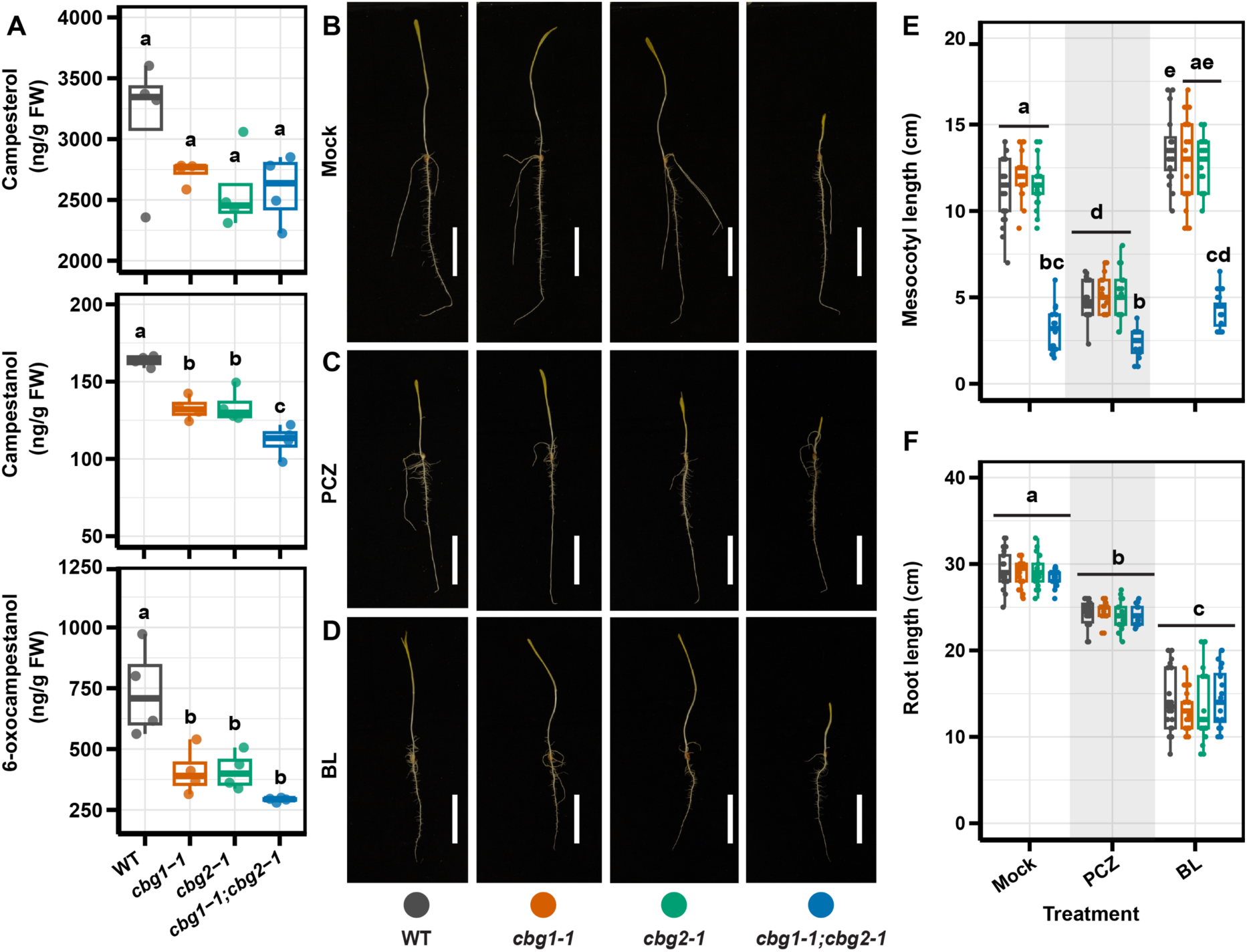
Effects of *Cbg1 and Cbg2* mutations on BR metabolite levels and seedling response to BL and PCZ. **(A)** BR metabolite levels of campesterol, campestanol and 6-oxocampestanol in 21-day-old seedlings (n=4), expressed as ng/g fresh weight (FW). **(B-D)** Representative whole seedlings of WT, *cbg1-1*, *cbg2-1*), and *cbg1;cbg2* double mutant (from right to left) grown for 5 days under **(B)** mock, **(C)** PCZ (200 µM), and **(D)** BL (2 µM) treatments. Scale bar = 8 cm. **(E)** Mesocotyl length and **(F)** Primary root length under the indicated treatments. Sample sizes (n) for mock, PCZ, and BL treatments, respectively were WT (23, 18, 20), *cbg1-1* (20, 21, 21), *cbg2-1* (21, 21, 21), and *cbg1-1;cbg2-1* (17, 17, 20). Boxplots show median, interquartile range, and individual biological replicates. Different lowercase letters indicate significant differences among genotypes based on one-way ANOVA for (A) and two-way ANOVA for (E-F), followed by Sidak adjusted multiple comparisons (*P < 0.05*). For (E-F), comparisons were performed across treatments.

### *cbg1*;*cbg2* double mutants phenocopy BR biosynthetic inhibition

To further test whether CBG1 and CBG2 function in BR biosynthesis, we chemically altered BR levels and quantified BR-responsive seedling traits, especially mesocotyl and root length (**Figure 3B-3D**). Because plant growth defects caused by BR deficiency in maize can be restored with exogenous BL (Makarevitch et al. 2012), the end product of the BR biosynthetic pathway, we tested whether BL treatment could rescue growth defects in the *cbg* mutants. In parallel, we used the BR biosynthetic inhibitor Propiconazole (PCZ) to assess whether chemically induced BR deficiency phenocopies the growth defects observed in the *cbg* double mutants. PCZ inhibits BL (and/or CS) biosynthesis by inhibiting activity of the CYP90C1 and CYP90D1 enzymes (Hartwig et al. 2012; Oh et al. 2016), which are downstream of CYP90A1 in the biosynthetic pathway. Wild-type, single mutant (*cbg1-1* and *cbg2-1)* and double mutant (*cbg1-1*;*cbg2-1*) seedlings were grown in darkness for 6 days under mock, PCZ, or BL treatment. Mesocotyl and root elongation were quantified as described (Hartwig et al. 2012; Zebosi 2024).

Under mock conditions in the dark, single mutant mesocotyl lengths (*cbg1-1* or *cbg2-1)* were indistinguishable from that of their wild-type siblings, whereas double mutants (*cbg1;cbg2*) exhibited significantly shorter mesocotyls (**Figure 3B** and **3E**), corresponding to approximately 72.4% reduction relative to the wild type (*p* < 0.05). In the absence of light, the impaired mesocotyl etiolation observed in the double mutants represents a typical skotomorphogenic phenotype associated with BR deficient mutants in maize, rice, and Arabidopsis (Mori et al. 2002; Hong et al. 2005; Hartwig et al. 2011; Makarevitch et al. 2012; Best et al. 2016). Also consistent with the hypothesis that *cbg1;cbg2* double mutants are BR-deficient, but the single mutants are not, PCZ treated wild-type and single mutants (*cbg1-1* and *cbg2-1)* showed significantly shorter mesocotyl length, resembling untreated *cbg1;cbg2* double mutants. In contrast, double mutants were less sensitive to PCZ treatment, as mesocotyl length did not differ between mock and PCZ treated double mutants (*p* > 0.05), consistent with already impaired BR biosynthesis (**Figure 3C** and **3E**). Application of exogenous, 2 µM BL did not significantly alter the elongation of the mesocotyl across genotypes (wild type, single mutants, and double mutants) relative to the mock treated (*p* > 0.05, **Figure 3C** and **3E**), indicating that BL treatment alone was insufficient to restore mesocotyl elongation of the *cbg1;cbg2* double mutants.

Seedling roots responded differently to PCZ and BL treatment compared to mesocotyl elongation. inhibition of BR biosynthesis with PCZ treatment resulted in a moderate but significant reduction in root length across all genotypes compared with mock controls, indicating that root growth is sensitive to BR biosynthetic inhibition even in the absence of full *Cbg* gene function. In contrast, treatment with 2 µM BL strongly inhibited root elongation across all genotypes (**Figure 3F**) consistent with previous studies in Arabidopsis and rice that showed BRs can promote root elongation at low concentrations but inhibit it at higher concentrations (Müssig et al. 2003; Tong et al. 2014). The moderately high BL concentration used in this study may therefore account for the inhibition of root elongation observed across genotypes. Overall, the primary root length was significantly affected by treatment (*p* < 0.05) but not by genotype (*p* > 0.05), with no significant differences among all genotypes within treatment. These findings indicate that shoot and root tissues respond differently to altered BR levels and suggest that *Cbg1* and *Cbg2* may play a more prominent role in regulating shoot growth rather than root elongation under the conditions examined, consistent with their markedly higher gene expression levels in shoots than in roots (**Figure 1B**).

## Discussion

Among brassinosteroid biosynthetic enzymes, cytochrome P450 CYP90A/CPDs are monooxygenases that play fundamental roles in plant growth and development, as demonstrated by functions extensively characterized in rice and Arabidopsis (Ohnishi et al. 2012; Zhan et al. 2022). However, their developmental roles in maize remain uncharacterized (Zebosi et al. 2024). Here, we show that duplicate maize CYP90A genes, *Cbg1* and *Cbg2*, collectively regulate plant architecture through affecting BR biosynthesis, as loss of both genes together resulted in several vegetative and reproductive defects whereas single mutants remained phenotypically normal under our field and greenhouse conditions.

The absence of obvious developmental defects in either cbg1 or cbg2 single mutants indicates functional redundancy between these duplicate genes. Although the subgenome 1 homeolog (*Cbg1*) is expressed at much higher levels in most tissues (**Figure 1B**), loss of *Cbg1* alone does not result in obvious developmental defects. Similarly, we found that levels of the downstream metabolites campestanol and 6-oxocampestanol were similarly reduced in seedling shoots of either single mutant, again despite the expression differences between *Cbg1* and *Cbg2*. These findings may suggest that the expression of *Cbg2* is sufficient to maintain normal plant development in the absence of *Cbg1*, either at its normal expression levels or such as by feedback regulation that could, for example, induce *Cbg2* expression in *cbg1* mutants. Regardless, the data indicate both genes must be disrupted before BR biosynthesis is sufficiently impaired to affect normal plant growth and development.

The spectrum of developmental defects observed in *cbg1;cbg2* double mutants is consistent with impaired BR biosynthesis (Makarevitch et al. 2012). As for mutants of single-gene steps in BR biosynthesis in maize such as *brd1*, *nana1*, and *nana2* (Hartwig et al. 2011; Makarevitch et al. 2012; Best et al. 2016; Castorina et al. 2018), *cbg1;cbg2* double mutants exhibit a similar, severe dwarf stature with compressed internodes that likely reflects reduced cell elongation (Kir et al. 2015). Similarly, erect leaves and an enlarged auricle phenotype are observed in *cbg1;cbg2* mutants, a hallmark of BR deficiency in rice, barley, and maize (Yamamuro et al. 2000; Chono et al. 2003; Kir et al. 2015; Best et al. 2016), further supporting the interpretation of impaired BR function. It will be interesting to determine if the consistent “cabbage head” phenotype we observed is common to other BR biosynthetic mutants if they are converged into the W22 or other, appropriate inbred genetic backgrounds. BR deficient mutants in maize often exhibit aberrant sex determination (Makarevitch et al. 2012; Best et al. 2016), commonly described as tassel feminization due to the formation (persistence) of silks in the tassels; however, this phenotype was not observed in *cbg1;cbg2* double mutants. The lack of silks in the tassel may suggest not all disruptions in BR biosynthesis produce equivalent effects on sex determination, or could again be related to differences in genetic background between the lines in our studies and the range of backgrounds used in other studies, or to environmental effects. We characterized the *cbg1;cbg2* double mutants in the W22 inbred, whereas *brd1* showed feminization in the B73 inbred (Makarevitch et al. 2012). On the other hand, for double mutants of the CYP90C1/CYP90D1 step in BL biosynthesis grown under our field conditions, the B73 genetic background completely suppressed silk formation in the tassel while others were permissive (Zebosi 2024). These results are consistent with natural genetic modifier loci that can suppress or enhance this phenotype.

Our metabolic profiling results qualitatively support that *Cbg1* and *Cbg2* function in BR biosynthesis, but comprehensive profiling of BR metabolites will be required to precisely demonstrate their biochemical functions within the pathway. It is unknown how much bio-active BRs are produced in the *cbg* mutants as their level was below detection level, even in the wild-type samples. It is interesting that downstream intermediates, such as campestanol and 6-oxcampestanol, were reduced in *cbg1;cbg2* double mutants as the *Atrot3/Atcyp90d1* double mutant were shown to have no difference in accumulation of these BR metabolites (Kim et al. 1998). This may suggest that as compared to Arabidopsis, for example, maize differs in how the BR pathway is regulated, in feedback mechanism and/or in the overall pool of p450 enzymes that can potentially compensate for loss of CPD.

In addition to developmental defects and metabolite measurements, pharmacological assays further support impaired BR biosynthesis In *cbg1;cbg2* mutants. Propiconazole (PCZ) treatment phenocopied the reduced mesocotyl elongation observed in *cbg1;cbg2* mutants, and in turn the double mutants showed reduced sensitivity to PCZ, two findings consistent with compromised BR biosynthesis (Hartwig et al. 2011). Reduced mesocotyl elongation under mock conditions in the dark reflects impaired skotomorphogenesis, a characteristic of BR deficiency across plant species (Hong et al. 2002). In contrast, exogenous brassinolide (BL) failed to rescue mesocotyl elongation in *cbg1;cbg2* double mutants although biological activity in our treatments was confirmed by BL’s inhibition of root elongation. BR deficient mutants in Arabidopsis can be restored by exogenous BL (Clouse 2002). In maize, exogenous BL partially rescues the *brd1* mutant, which is defective in late steps of BR biosynthesis including the final C-6 oxidation (Makarevitch et al. 2012). However, comparable experiments have not been reported for the earlier BR biosynthetic mutants such as *na1* and *na2* (Hartwig et al. 2011; Best et al. 2016), which act early in the biosynthetic pathway just up- and downstream of where *Cbg* genes function. Thus, the inability of BL to rescue *cbg1*;*cbg2* mutants does not imply that maize cannot respond to exogenous BL but instead demonstrates that BL rescue varies among BR biosynthetic mutants. Studies in rice and *Brachypodium distachyon* provide additional context on BR biosynthesis in grasses for interpreting the lack of BL rescue in *cbg1;cbg2*. In rice, exogenous BL partially rescues BR deficient mutants (Hong et al. 2002; Mori et al. 2002; Hong et al. 2003) yet in biochemical studies in rice and *Brachypodium distachyon* BL has not been detected, supporting a hypothesis that BL’s precursor castasterone (CS) is the bio-active BR in these species (Kim et al. 2008; Roh et al. 2020). Whether maize BR bio-activity acts similarly remains unclear. On the other hand, because exogenous BL rescues some BR deficient mutants in both maize and rice, the lack of rescue in *cbg1;cbg2* double mutant mesocotyls cannot be explained simply by the use of CS rather than BL as the predominant endogenous brassinosteroid. Further studies including more comprehensive quantification of BR intermediates and feeding experiments with CS, will help determine why exogenous BL fails to restore mesocotyl elongation in *cbg1;cbg2* double mutants.

At the seedling stage, *cbg1;cbg2* double mutants grown under either greenhouse or growth chamber conditions exhibited severe shoot growth defects but no obvious defects in root development, suggesting that *Cbg1* and *Cbg2* play a prominent role in shoot than root development. This observation is consistent with higher expression of *Cbg1* and *Cbg2* in shoots compared to roots (**Figure 1B**). In addition, the preferential inhibition of root elongation by exogenous BL, suggests that shoot and root tissues differ in their sensitivity to BR levels, consistent with studies in Arabidopsis showing that optimal BR concentrations vary among tissues and developmental zones, particularly within the root where local BR biosynthesis is tightly regulated (Müssig et al. 2003; Singh and Savaldi-Goldstein 2015; Vukašinović et al. 2021).

Although a few BR biosynthetic mutants, including *na1*, *na2*, *m30,* and *brd1* have been characterized in maize (Hartwig et al. 2011; Makarevitch et al. 2012; Best et al. 2016; Sun et al. 2024), several steps in the grid-like BR biosynthetic pathway, including C-22 hydroxylation, C-23 hydroxylation, and C-3 oxidation reactions remain genetically uncharacterized (Zebosi 2024; Zebosi et al. 2024). Our characterization of *Cbg1* and *Cbg2* addresses one of these gaps by showing that two maize CYP90A orthologs function redundantly to regulate BR biosynthesis and plant development. The severe pleiotropic defects of *cbg1;cbg2* double mutants indicate complete loss-of-function of *CBG* genes is detrimental to plant fitness but raise the possibility that fine-tuning of *CBG1* and *CBG2* expression may provide an effective strategy to optimize plant architectural traits such as plant height and leaf angle without compromising other important agronomic traits.

## Materials and methods

### Genetic stocks and mutant isolation

UniformMu insertion alleles in *cbg1* (Zm00001eb196530) and *cbg2* (Zm00001eb093140) were obtained from the Maize Genetics Cooperation Stock Center (McCarty et al. 2005; Woodhouse et al. 2025). The *cbg1* alleles included *cbg1-1* (UFMu-10570; *mu1086371*), *cbg1-2* (UFMu-04183; *mu1025370*), and *cbg1-3* (UFMu-04179; *mu1038572*), whereas *cbg2* alleles included *cbg2-1* (UFMu-06432; *mu1047171*) and *cbg2-2* (UFMu-08204; *mu1059083*).

Individual plants heterozygous for *cbg1* and *cbg2* alleles were identified and self-pollinated to make a segregating population. Because none of the single mutants displayed obvious developmental defects, *cbg1* and *cbg2* double mutants were generated to test the functional redundancy between the two genes. Plants homozygous for *cbg1* (*cbg1-1*, *cbg1-2* or *cbg1-3*) allele were crossed to *cbg2* (*cbg2-1* or *cbg2-2*) alleles in the following combinations: *cbg1-1* x *cbg2-1*, *cbg1-2* x *cbg2-1*, *cbg1-3* x *cbg2-1* and *cbg1-1* x *cbg2-2*. The resulting F1 plants were self-pollinated to yield F2 populations, in which *cbg1;cbg2* double mutants segregated at an expected frequency of 1/16. The F2 were phenotyped for plant height, tassel branch number, leaf size and leaf angle.

F2 plants homozygous for *cbg1* and heterozygous for *cbg2* (*cbg1-1; cbg2-1/+*, *cbg1-2; cbg2-1/+*, *cbg1-3; cbg2-1/+*, and *cbg1-1; cbg2-2/+*) were self-pollinated to create F3 populations, in which double mutants segregated at frequency of 1/4. In addition, allelism tests were performed by crossing *cbg1-1/+;cbg2-1* with *cbg1-2/+;cbg2-1*, *cbg1-3/+;cbg2-1* and *cbg1-1/+;cbg2-2*. Chi-square tests were performed to evaluate whether observed progeny segregation ratios differed from the expected Mendelian ratios.

### Growth conditions

Field-grown plants were grown in the summer maize nursery at Curtiss Farms, Iowa State University in Ames, Iowa. For greenhouse experiments, plants were grown in 1-gallon pots under standard conditions with a 16h light / 8 hr dark cycle under supplemental lighting and temperature of 27°C during the day and 21°C at night (Zebosi et al. 2026).

### Genotyping

To confirm *Mu* insertions in *cbg1* and *cbg2*, kernels from each stock were planted in the field, and genomic DNA was extracted from pools of 10-20 plants using the guanidine-hydrochloride method (Gao et al., 2010; Zebosi et al., 2024). The *Mu* insertion sites were verified using either a four-primer or three-primer PCR approach and the size of amplified fragments in gel electrophoresis. PCR was performed as described by (Zebosi et al. 2025), using either two Mu-specific and two gene-specific primers or two *Mu*-specific primers and one gene-specific primer (**Supplemental Table 1**). The PCR amplicons were visualized on 2% agarose gels. Pools containing the expected *Mu* insertions were deconvoluted to individual plants which were genotyped by the same assays described above.

### Phylogenetic analysis

Brassinosteroid Cabbage/CYP90A protein sequence from Arabidopsis, maize, rice, Brachypodium, Medicago, Sorghum, Setaria, soybean, Solanum lycopersicum, and Physcomitrella were identified by BLAST searches from Phytozome (Goodstein et al. 2012) as previously described (Zebosi et al. 2024). Protein sequences were aligned using the MAFFT multiple sequence alignment program (Katoh et al. 2002), and the resulting alignment was used to construct a maximum-likelihood phylogenetic tree of the CYP90A gene family using FastTree (Price et al. 2010). The resulting Newick file was visualized and interpreted using Dendroscope (Huson et al. 2007). Bootstrap analysis was performed with 1000 replicates, and bootstrap values were added to each node to show the strength of the branch. The phylogenetic tree was rooted using Physcomitrella genes as out-groups.

### Metabolite measurements

For metabolite measurement, shoot tissues were harvested wild-type, *cbg1-1*, *cbg2-1,* and *cbg1-1;cbg2-1* double mutant seedlings. Plants were grown for 24 days in the greenhouse, cut off at the soil line and trimmed of expanded leaves to retain a 3 cm segment of shoot. Four biological replicates were collected per genotype, with each replicate consisting of pooled tissue from eight plants. Samples were immediately frozen in liquid nitrogen and were then processed and metabolites quantified as described by (Dilkes and Best 2025).

### PCZ and BL feeding experiments

Seeds of wild-type, *cbg1-1*, *cbg2-1* and double mutants (*cbg1-1;cbg2-1*) were planted in germination paper soaked in water, 200uM propiconazole (PCZ) and 2uM brassinolide (BL) solution, rolled up and placed in buckets as previously described (Draves et al. 2022). Seedlings were grown under dark conditions at 28°C and 5 days after planting mesocotyl and primary root length were measured.

### Accession numbers

*Cabbage1* (*Cbg1*) : GRMZM2G012391 (B73 RefGen_v3), Zm00001d052475 (Zm-B73-REFERENCE GRAMENE-4.0), Zm00001eb196530 (Zm-B73-REFERENCE-NAM-5.0) *Cabbage2* (*Cbg2*) : GRMZM2G162737 (B73 RefGen_v3), Zm00001d004957 (Zm-B73-REFERENCE GRAMENE-4.0), Zm00001eb093140 (Zm-B73-REFERENCE-NAM-5.0)

## Acknowledgments

We would like to acknowledge the assistance of ISU staff at Curtiss Farms and the Molecular Biology Building greenhouses. Thanks to the Maize Genetics Cooperation Stock Center for generously providing maize seeds. This work was supported by the NSF Plant Genome Research Program (Award #1238202) and Hatch project IOW05745 from the Iowa Agriculture and Home Economics Experiment Station at Iowa State University, awarded to E.V.

## Author contributions

B.Z. and E.V. designed the research, B.Z, J.S., E.U.W, N.B. and E.V. performed experiments and analyzed data. B.Z., N.B. and E.V. wrote the manuscript. E.V. acquired funding. All authors edited and approved the manuscript.

