## supplemental figures for "*Cabbage1* and *Cabbage2* encode CYP90A orthologs that function redundantly to regulate plant development in maize"

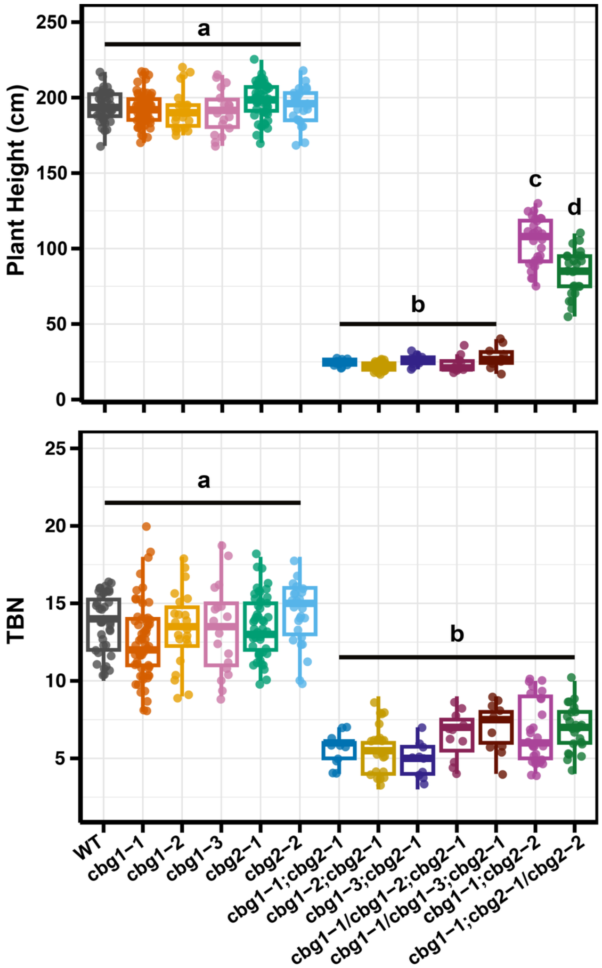


**Supplemental Figure 1:** Plant height and tassel branch number (TBN) across field-grown genotypes. Data were collected from the same experiment as in Figure 1 but single mutants are not shown there. Plant height is reported in centimeters (cm) and TBN as the number of tassel branches. Sample sizes (n) in the order shown on the x-axis were 40, 62, 22, 50, 25, 18, 10, 24, 10, 11, 10, 28, and 25. Different lowercase letters indicate significant differences among genotypes (ANOVA followed by a post hoc multiple comparisons test with $P < 0.05$).

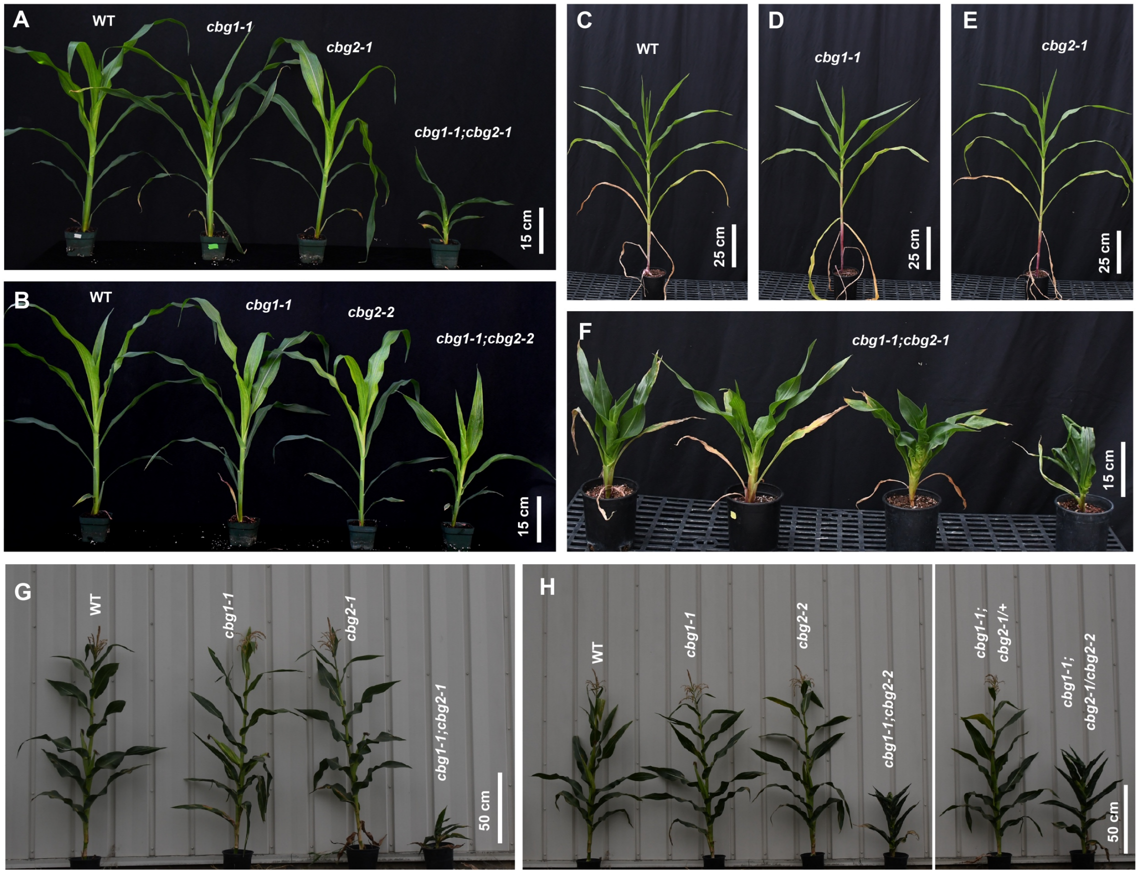


**Supplemental Figure 2: phenotypes of *cabbage* (*cbg*) mutants.**
**(A-B)** Plants in the W22 inbred background grown under greenhouse conditions. **(A)** Wild-type (WT) and single mutants (*cbg1-1* and *cbg2-1*) show no obvious phenotypic differences, whereas the *cbg1-1;cbg2-1* double mutant (left) displays dwarf stature and rolled leaves. **(B)** Similar to **(A),** but with the *cbg2-2* allele. The *cbg1-1;cbg2-2* double mutant exhibits a milder semi-dwarf phenotype than the *cbg1-1;cbg2-1* double mutant, while the single mutants resemble WT. **(C-F)** Plants after one generation of crossing to B73 inbred background (X1)**.** The *cbg1-1;cbg2-1 d*ouble mutant exhibits dwarf stature with variable expressivity (**F**), whereas the corresponding single mutants are phenotypically indistinguishable from WT. **(G-H)** Mature field grown plants in the W22 background. **(G)** The same genotypes as shown in (A), showing the *cbg1-1;cbg2-1* double mutants under field conditions. **(H)** The same genotypes as in (B), including *cbg2-1*/*cbg2-2* compound heterozygotes.

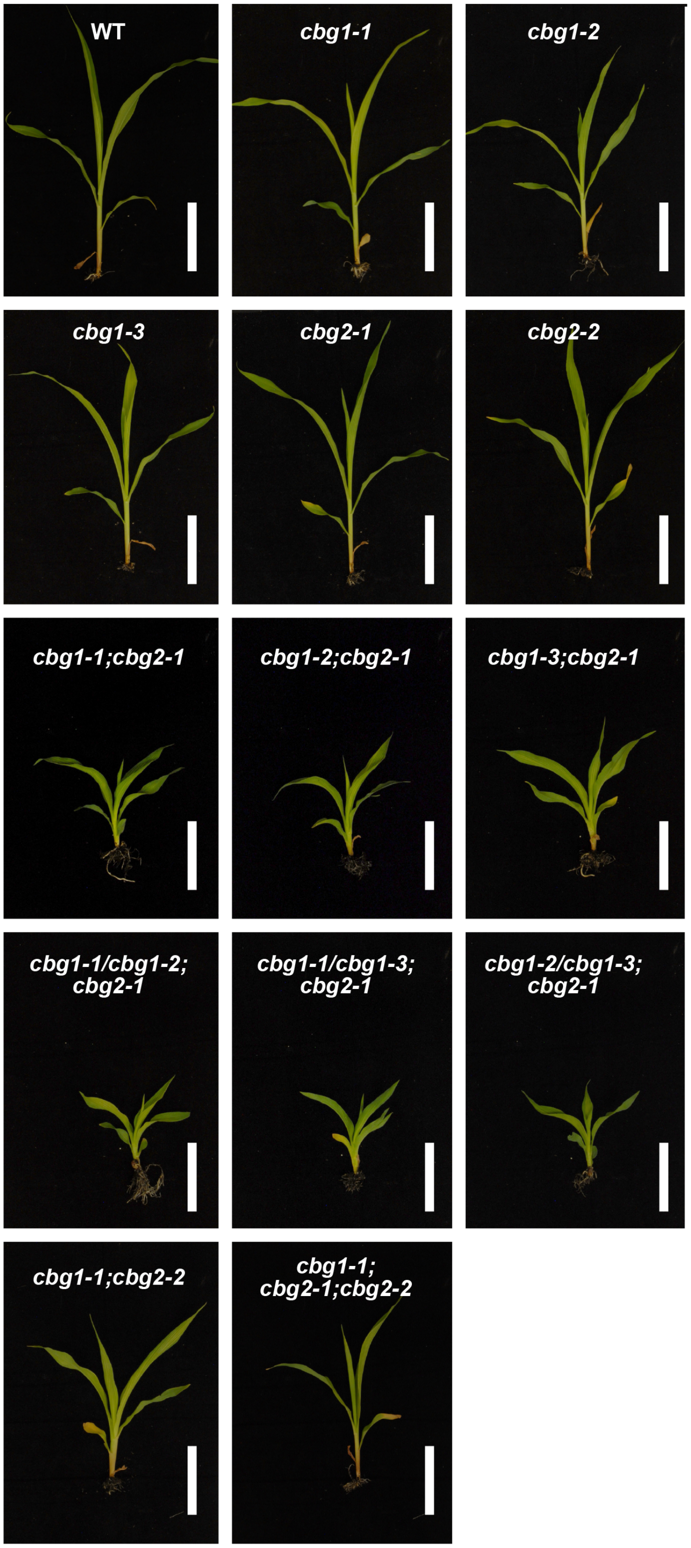


**Supplemental Figure 3: Seedling phenotypes of cbg single mutants, double mutants, and compound heterozygotes.** Three-week-old seedlings in W22 genetic background grown in greenhouse conditions. Scale bar = 8 cm

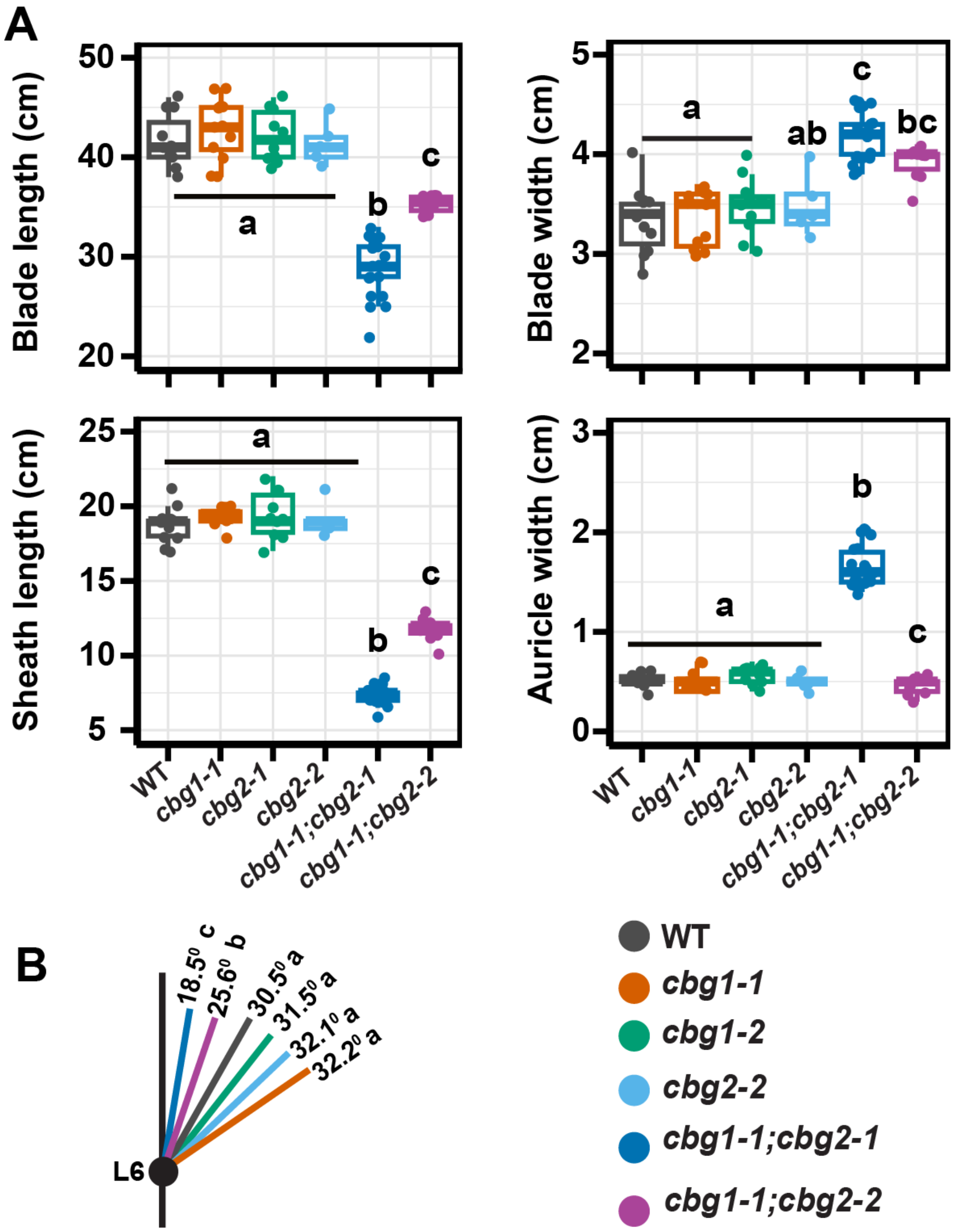


**Supplemental Figure 4: Leaf morphology of greenhouse-grown *cbg* mutants.**
**(A)** Blade length, blade width, sheath length, and auricle width of leaf 6 measured across genotypes; WT (n=11), *cbg1-1* (n=12), *cbg1-2* (n = 10), *cbg2-2* (n = 6), *cbg1-1;cbg2-1* (21) and *cbg1-1;cbg2-2* (n = 10). Boxplots show the median, low quartile, upper quartile with whiskers extending 1.5 x the interquartile range. Individual points represent biological replicates. **(B)** Schematic representation of leaf angle at leaf 6 of mean leaf angle for WT (n=11), *cbg1-1* (n=7), *cbg1-2* (n = 10), *cbg2-2* (n = 12), *cbg1-1;cbg2-1* (n = 10) and *cbg1-1;cbg2-2* (n = 8). Colored circles correspond to the genotype shown in the legend. Different lowercase letters indicate significant differences among genotypes based on one-way ANOVA followed by a multiple comparison test ( $P < 0.05$).

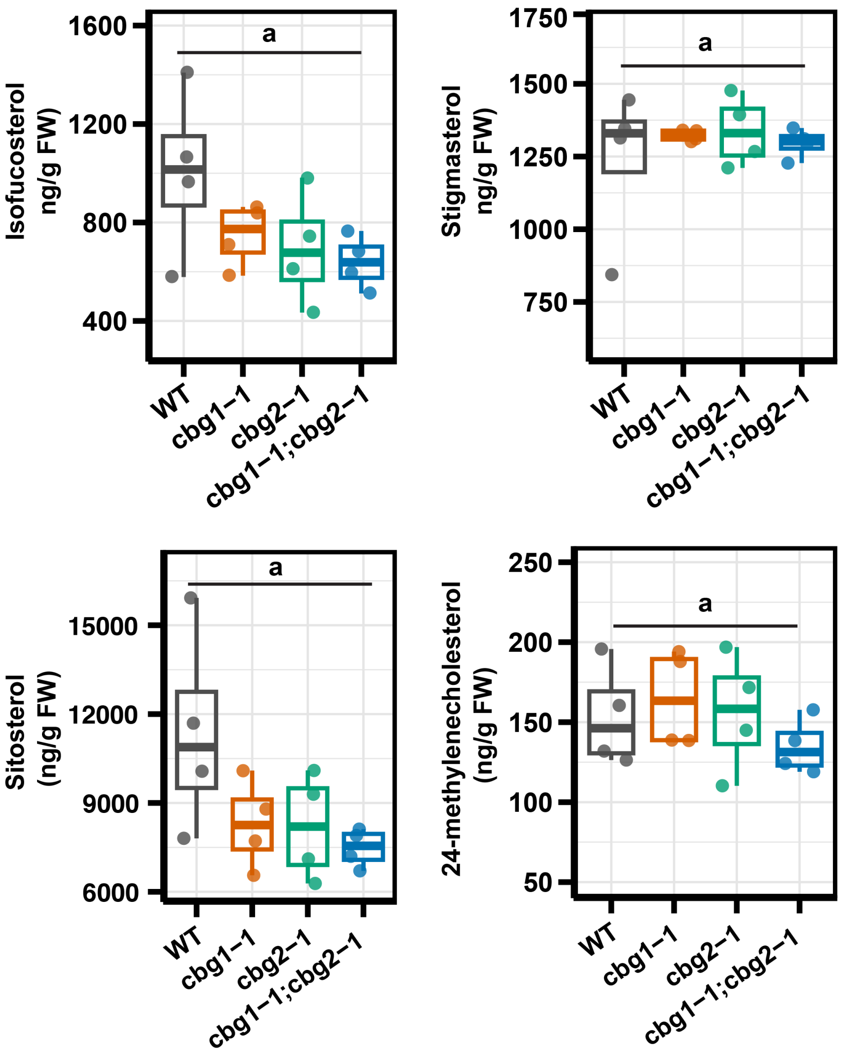


**Supplemental Figure 5: Metabolite profiling**

BR metabolite levels in 21-day-old seedlings (n=4). Different lowercase letters indicate significant differences among genotypes based on one-way ANOVA followed by Sidak adjusted multiple comparisons ($P < 0.05$).

**Supplemental Table 1.** Primers used in this study

| Primer | Sequence | Genotyping details |
|---|---|---|
| BZ600 | CACACGGTGAGGTTGAAGGTGA | *cbg1-1* |
| BZ597 | GCAACAGCGCATCACTCACAC | |
| Mu1 | GCCTCCATTTCGTCGAATCCC | |
| Mu2 | GCCTCTATTTCGTCGAATCCG | |
| EUO 1416 | GATGTACCTTGAGCTGCGCCAG | *cbg1-2* and *cbg1-3* |
| EUO 1417 | GAGTACGTGAAGCTCGTCGACG | |
| Mu1 | GCCTCCATTTCGTCGAATCCC | |
| Mu2 | GCCTCTATTTCGTCGAATCCG | |
| BZ626 | CCACTATGACGCTGCTACTATAATAAGG | *cbg2-1* and *cbg2-2* |
| BZ634 | CCGCAGCCAACCAACCAACC | |
| Mu9242 | AGAGAAGCCAACGCCAWCGCCTCYATTTCGTC | |
